# Climate change, infectious disease, and the spread of microblades across the Qinling-Huaihe line

**DOI:** 10.64898/2026.09.14.751440

**Authors:** Shoshana Elgart, Kenichi Aoki, Marcus W. Feldman

## Abstract

Aoki et al. (2023) proposed that the sharp difference in microblade distribution between the north and south of the Qinling-Huaihe (H-Q) line before the early Holocene could have been due to a higher frequency of pathogens in the south. Greater susceptibility to pathogen-induced disease among northerners could have effectively prevented migration across the H-Q line. Here, we explore the possibility that migration between the north and south could have been stalled by a vector-borne disease whose vector distribution was climate dependent. The original wave equation approach is extended to include climate variation in the forms of (i) a period of linear warming; (ii) a period of linear cooling; and (iii) a period of temperature oscillations. Simulations of this extended model are carried out using published estimates of the climate record for the Northern Hemisphere since the Upper Paleolithic. This analysis suggests possible time intervals during which microblades could have reached the south.

## 1 Introduction

A major feature of the Upper Paleolithic, from approximately 50 to 12 kya, is accelerated differentiation in human tool-making, from blade construction to the fashioning of scrapers, chisels, and bone awls (Bar-Yosef, 2002; Derevianko, 2010). Small pointed microblades, acquired via pressure-flaking techniques, appeared in northern and north-east Asia during the Upper Paleolithic, between 40-25 kya (Gómez-Coutouly, 2018; Yi et al., 2016). It has been suggested that microblades facilitated the hunting of large mammals in the so-called Mammoth Steppe, the production of fitted warm clothing from furs and skins, and the development of composite tools such as sickles (Graf, 2010; Kato, 2014; Yi et al., 2016; Liu et al., 2026).

Microblades dating to the Upper Paleolithic are frequent in northern and northeast China, but essentially absent at this time below the Qinling–Huaihe (H-Q) line that divides northern and southern China (Bar-Yosef, 2002; Qu et al., 2013). This has suggested the existence of an informal *boundary* that restricted the range expansion of microblade-using foragers in northern China to territories *north of* the H-Q line, limiting their interaction with southern-based communities (Aoki et al., 2023; Yi et al., 2016).

Aoki et al. (2023) suggest that increased pathogen spread in the tropical and sub-tropical regions of southern China—particularly endemic sources of disease such as *Plasmodium vivax* malaria—may have played substantial roles in the formation of this informal border. They propose two reaction-diffusion models of northern and southern-forager migration in the presence of *southern-only* pathogens to which northerners are susceptible. The first is a “minimal” two-population model in which no distinction is made between northern foragers with and without microblades, while the second is a three-population model, in which microblade-possessing communities of northerners are treated as a separate group, and cultural transmission with conformity affects the rates at which non-microblade possessing northerners acquire microblades. They obtain conditions for the existence of a *stationary front* between the density of northerners with microblades and southerners without microblades.

Here, we incorporate the effect of climate variation on both disease burden in the southern population and the cultural transmission of microblade use into the model of Aoki et al. (2023). In particular, the late Upper Paleolithic overlaps with the Last Glacial Maximum (LGM, 27-19 kya), which led to significant fluctuations in temperature and humidity on a centuries-long scale (Kravchinsky et al., 2021). Recent literature (e.g., Kravchinsky et al. (2021); Zhang et al. (2025)) has used fossil pollen records to argue that climate in northern and southern East Asia varied dramatically in the millennia associated with the LGM, with regions south of the H-Q line remaining substantially warmer and wetter than those north of it. The millennia preceding the LGM, moreover, were distinguished by a number of *quasi-periodic* warming and cooling cycles known as Dansgaard-Oeschger (DO) events (Li et al., 2017); in particular, the first through eighth of these events likely took place *after* the arrival of microblades in northern China (38-27 kya).

This climate variation would have driven changes in disease burden across north and south China. As discussed by Aoki *et al*., vector-borne diseases such as *Plasmodium vivax* malaria may have limited migration of northern groups into southern territories. The density of the *Anopheles sinensis* mosquitoes, those chiefly responsible for malaria transmission in China, is strongly climate-dependent. For example, the *A. sinensis* egg-laying rate depends on the availability of *standing water* for breeding, which increases when humidity is high and falls during a glacial period (Brown et al., 2023). *A. sinensis* hatching rates are reduced as temperature decreases (from under 0.34 days*^−^*^1^ at 25–30*^◦^* C, to 0.15 days*^−^*^1^ at 22*^◦^* C), and adult mosquitoes are known to enter diapause below 10*^◦^* C (Feng et al., 2017). *A. sinensis* individuals do not generally survive extreme temperatures, below 0*^◦^* C and above 40*^◦^* C (Rossati et al., 2016; Feng et al., 2017).

This temperature dependence of *Anopheles* can be compared to estimates of paleo-climate around the time of the LGM, which is typically inferred by measuring *δ*^18^*O isotope levels* in ice cores (Andersen et al., 2004; Liu et al., 2023). In particular, the relationship between changes in temperature and *δ*^18^*O* isotope levels has been generally assumed to be linear or quadratic (Gaskell and Hull, 2023; Liu et al., 2023). For instance, each unit difference Δ*δ*^18^*O* in isotopic values at the Huascarán ice-cores and the corresponding change Δ*T* in local temperature have been estimated to satisfy

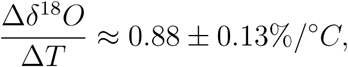

if the local temperature change Δ*T* scales linearly with the *global* mean surface temperature (GMST) (Liu et al., 2023, pg. 6).

Assuming this linear approximation for *δ*^18^*O* as a function of GMST, and an increase of Δ*δ*^18^*O ≈* 6.5% in Huascaŕan isotopic values from the LGM to the pre-industrial age, Liu et al. (2023) estimate the LGM-to-pre-industrial change in GMST at 5.9 *±* 1.2*^◦^*C. Assuming the same temperature change in East Asia, the mean LGM temperature around Beijing in north China could be estimated at *≈* 5.6*^◦^*C (using an approximate pre-industrial temperature of 11.5*^◦^* C in Beijing (Ren et al., 2022)). Similarly, using an estimated pre-industrial temperature of 16*^◦^ −* 22*^◦^* C in southern Lake Poyang (Miao et al., 2025), the mean LGM temperature in south China could be estimated at 10.1*^◦^ −*16.1*^◦^*C. Therefore, since *Anopheles sinensis* mosquitoes typically enter diapause below 10*^◦^* C, and rarely hatch below 16*^◦^* C (Feng et al., 2017), the ice core estimates of Andersen et al. (2004) suggest that vector populations could have been markedly reduced in south China (and negligible in north China) around the time of the LGM. At the time when microblades spread through north China, mosquito population sizes may therefore have differed substantially between warming and cooling events, and between northern and southern China. Variation in mosquito density would in turn have affected temporal and spatial variation in malarial prevalence (Rossati et al., 2016). Additionally, the possible use of microblades in the production of warm clothing, and the likely involvement of microblades in efficient hunting techniques (Elston and Brantingham, 2002; Yi et al., 2016), suggests that microblade possession may have given a fitness benefit to northern foragers during cooling events.

In Section 2, we formulate an extension to the microblade diffusion model of Aoki et al. (2023), under the assumptions that disease burden for northern populations is driven by a vector-borne disease and selection for microblade use is dependent on a single climate variable, generally assumed to be temperature. For simplicity, we refer below to the vector-borne disease as *P. vivax* malaria, and to the vector as *Anopheles* mosquitoes, but our models could apply more generally to vector-borne disease.

Our model involves explicit equations for temperature-dependent mosquito populations that transmit *P. vivax* in southern China, and introduces *climate-driven* selection into the cultural transmission of microblades between northern foraging communities. We assume that these two sets of climate-driven effects are *coupled*, in that higher fitness benefits for microblade possession correlate (under lower temperatures) with decreases in mosquito birth rates. Moreover, unlike in Aoki et al. (2023), we assume that southerners may convert to microblade use if there is sufficient contact with microblade-possessing northerners, possibly at slower rates than those associated with cultural transmission *among* northerners.

We simulate this model using temperature data inferred from *δ*^18^*O* climate records from the North Greenland Ice Core Project (NGRIP) to represent the period 35-7 kya (Andersen et al., 2004). Possible *time windows* during which southerners adopt microblade use, as functions of the human travel rate, the disease burden in northerners, and the rate of north-south cultural transmission are analyzed.

## 2 When southerners adopt microblades: simulations using climate data from the North Greenland Ice Core Project (NGRIP)

We first extend the two-population model of Aoki et al. (2023) to incorporate a disease transmitted by climate-dependent *vector* populations, which introduces seasonal selection to the cultural transmission of microblades between northerners. In particular, each mosquito population is assumed to be localized in space, following Feng et al. (2017), who conclude that the ranges of *Anopheles* mosquitoes in China are generally limited to a kilometer. In contrast, the *demography* of the mosquito populations is assumed to vary spatio-temporally with a single climate variable, assumed here to be temperature.

To model mosquito demography, the time-range of the two-population model equations (eqs. (2.1a)-(2.1b) in Aoki et al. (2023)) is restricted to a long fixed interval *T* := [*T*_0_*, T*_max_], associated with a sequence of *climate events* that force temperature changes across *T*. Also, let *D* denote a one-dimensional (south to north) spatial domain across which migration is assumed to occur, where (with appropriate non-dimensionalization) *D* is assumed to be the unit interval [0, 1]. Then define a *temperature function*

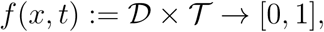

which measures a correlate of temperature at some time *t ∈ T* and point *x ∈ D* (where *x* = 0 and *x* = 1 at the extreme south and north of the spatial domain, respectively). It is assumed that *f* is continuous on the spatiotemporal domain *D×T* and monotonically decreasing in *x* when *t* = 0. At each point (*x, t*) *∈ D × T*, the mosquito birth rate is assumed to increase *linearly* with the temperature function *f* (*x, t*), approximating the dependence of *Anopheles* hatching rates on temperature (Section 1).

Individuals who have malaria in northern groups are assumed to be immediately removed from the population, and to produce no secondary cases, while malaria-infected individuals in southern groups are treated as asymptomatic, and remain in the population. As a result, malaria-infected northerners are assumed to be unable to transmit the disease to a susceptible mosquito in their environment, having exited the infecting population before additional human-mosquito interaction, but southerners *are* able to reinfect surrounding mosquitoes. However, secondary *P. vivax* cases in southerners that arise from within-host effects (i.e., the activation of hypnozoites) are not included, as *P. vivax* malaria recurrences span much shorter periods than the (century-long) time-scale assumed here (White, 2011).

Let *N*_1_(*x, t*) and *N*_2_(*x, t*) denote the respective densities of southern and northern foragers at a point in the spatiotemporal domain *D × T*, and let *S*(*x, t*) and *I*(*x, t*) := *D×T →* [0, 1] denote the *susceptible* and *infectious* mosquitoes’ densities, respectively, at each point (*x, t*) *∈ D×T*. Then, recalling that the mosquito populations are assumed to be localized in space, we have

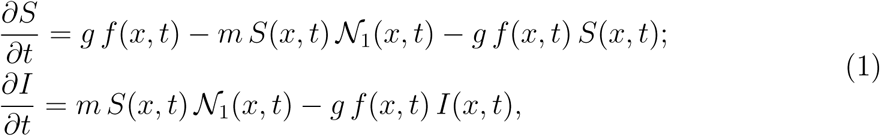

where the parameter *m* represents both the human-to-mosquito contact rate, and the probability of human-to-mosquito transmission following a single mosquito bite. The term *gf* (*x, t*) specifies the mosquito population growth rate at a point (*x, t*) *∈ D × T* for a fixed constant *g ∈* R^+^, which we treat as the *baseline* demography rate for mosquito populations. The total number of mosquitoes is constant over time, and (with appropriate non-dimensionalization) assumed to be 1.

Since southerners can eventually convert to microblade use, the southern population *N*_1_ is divided into a sub-population *C*_1_, representing southerners without microblades, and a sub-population *M*_1_ := *N*_1_ \ *C*_1_, of southerners with microblades. Therefore, we distinguish three populations of densities *C*_1_, *M*_1_, and *N*_2_, and denote their carrying capacities by *L*_1_, *K*_1_, and *K*_2_, respectively, and their intrinsic growth rates by *r_C_*_1_, *r_M_*_1_, and *r_N_*_2_, respectively. In the simplest case, it is generally assumed that the intrinsic growth rate of southerners with microblades equals that of northerners; whenever a value is assigned to *r_M_*_1_, it can always be taken to represent a value for *r_M_*_1_ = *r_N_*_2_.

Additionally, denote the spatial *diffusion rates* of the three human populations along the domain *D* by *δ_C_*_1_*, δ_M_*_1_, and *δ_N_*_2_, respectively. It is assumed in particular that *δ_M_*_1_ = *δ_N_*_2_ *> δ_C_*_1_ (following Yi et al. (2016), who suggest that microblade use may have extended the hunting range of a population). The Lotka-Volterra competition coefficients between the three populations are all assumed to equal a constant *b*.

Two directions of cultural transmission for microblade use are considered: *within* southern groups, and *between* northern and southern populations. In particular, the first process involves conformity bias: the rate at which individuals in *C*_1_ adopt microblade use after contact with individuals in *M*_1_ is supposed to be *ɛM*_1_*C*_1_*P*, where *ɛ ∈* R^+^ is a rate constant and

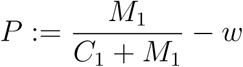

is positive when the proportion of microblade-possessing southerners exceeds the constant *w*, and negative otherwise. Further, we assume that the rate at which individuals in *C*_1_ adopt microblade use after contact with individuals in *N*_2_ is simply

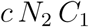

with a rate constant *c ∈* R^+^, so that microblade transmission between the northern and southern populations mimics an *epidemiological* process. Assume that *c < ɛ*, so that the influence of northerners on southerners’ microblade status is less than that of other southerners.

Microblade use dynamics in the southern population are also assumed to include *selection* for the cultural trait of microblade use, in which microblade non-possessors lose fitness as temperatures cool (following Yi et al. (2016), who propose that microblade use may have facilitated the production of warm clothing). The function *s*(*x, t*) *∈* [0, 1] denotes the loss of fitness in non-microblade possessing individuals (relative to microblade-possessing ones), which is assumed to depend exclusively on the temperature function *f* (*x, t*). In the simplest possible case, *s*(*x, t*) is set to be *V* exp (*−f* (*x, t*)) for a parameter *V ∈* [0, 1].

The full model system for *C*_1_, *M*_1_, and *N*_2_ then becomes:

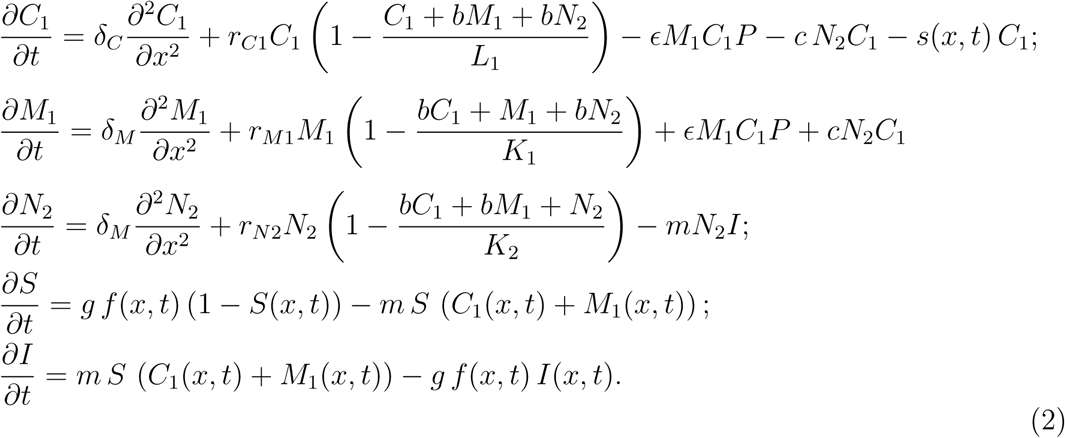

Here, as in eqs. (1), *S*(*x, t*) and *I*(*x, t*) denote the densities of susceptible and infectious vectors, respectively, at a point (*x, t*) in the spatiotemporal domain.

The initial and boundary conditions for System (2) are assumed to be of the form

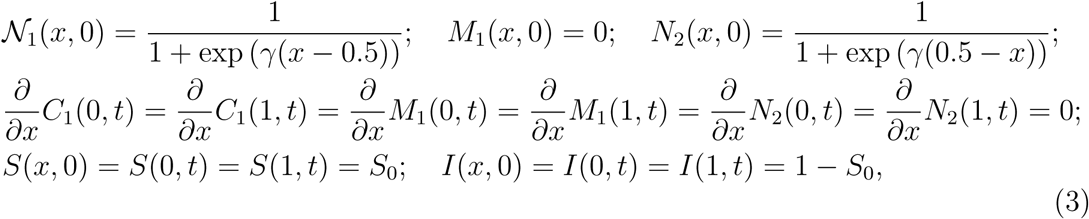

where the parameter *γ ≫* 1 is arbitrarily large. In particular, the initial distributions of northerners and southerners across *D* are modeled by sharp *sigmoid* curves with inflection points at *x* = 0.5. The density *N*_1_(*x,* 0) is near 1 on the whole interval [0, 0.5) and near 0 on the whole interval (0.5, 1], while the density *N*_2_(*x,* 0) := *N*_1_(1 *− x,* 0) satisfies the *reverse* conditions on the intervals [0, 0.5) and (0.5, 1].

The density of southerners possessing microblades, *M*_1_(*x, t*), is assumed to be zero for all *x* at *t* = 0. The boundary conditions for the human populations are assumed to be zero-flux, and the ratio of infectious-to-susceptible mosquitoes is initially fixed at a positive constant (1 *− S*_0_)*/S*_0_.

System (2) is simulated using climate data from the North Greenland Ice Core Project (NGRIP), which contains correlates of temperature in the Northern Hemisphere between 0-123 kya, recorded at 100-year intervals and measured using *δ*^18^*O* isotopes in two Greenland ice sheets (Andersen et al., 2004). In particular, the *δ*^18^*O* isotopic values in Andersen et al. (2004) are normalized to lie in the interval [0, 1], and substituted for the temperature function *f* (*x, t*). The simulation is initiated at 35 kya, corresponding to the approximate first observation of microblades in northern China (Yue et al., 2021), and continued until 7 kya, around the mid-Holocene.

Table 1 lists ranges for the parameter values used in the simulation of System (2), together with their sources where present. All parameters are in units of (1000 years)*^−^*^1^, referred to as *ky^−^*^1^, unless otherwise stated.

**Table 1:**
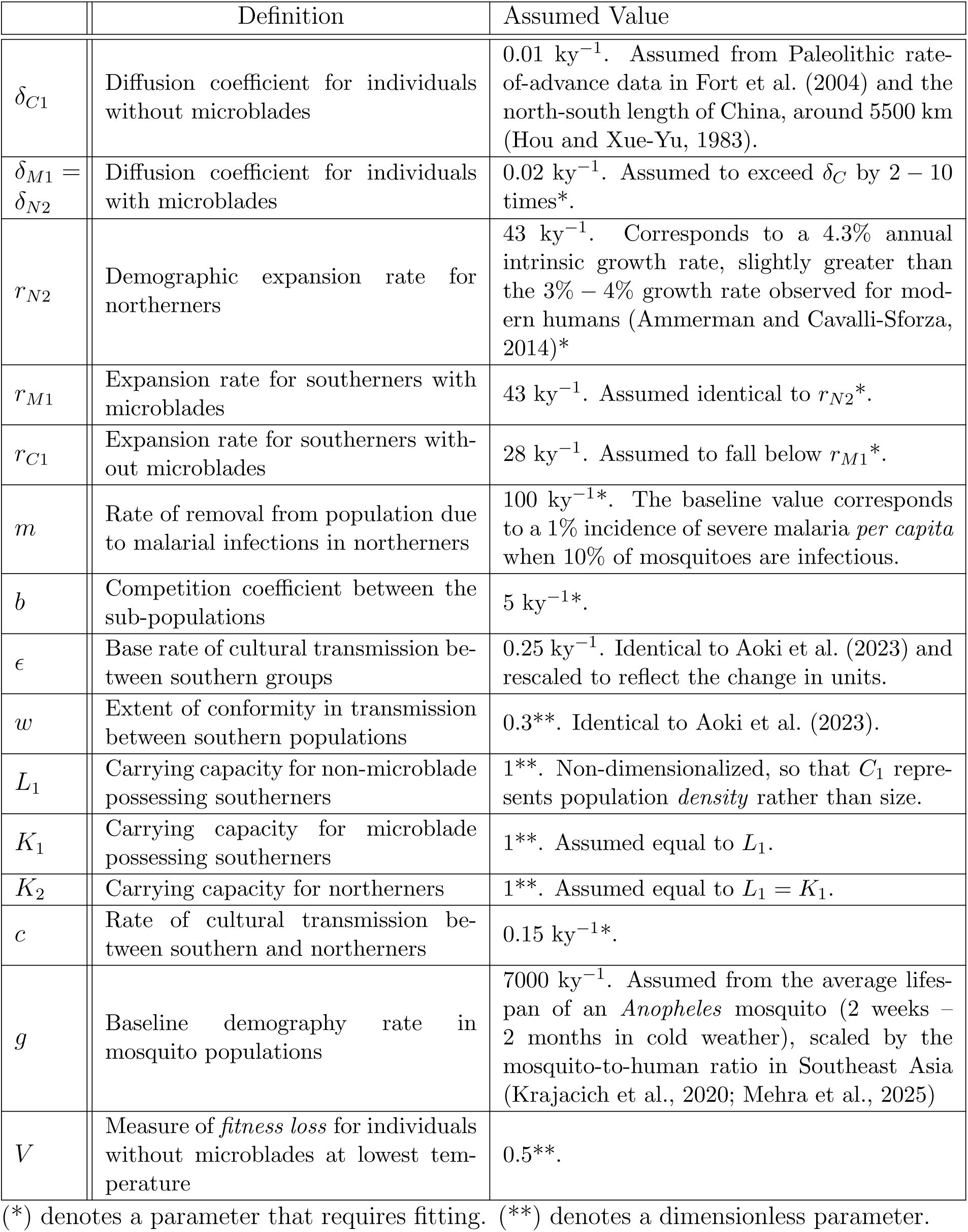
List of parameter values in the simulation of system (2).

|  | Definition | Assumed Value |
| --- | --- | --- |
| $\delta_{C1}$ | Diffusion coefficient for individuals without microblades | $0.01 \text{ ky}^{-1}$. Assumed from Paleolithic rate-of-advance data in Fort et al. (2004) and the north-south length of China, around 5500 km (Hou and Xue-Yu, 1983). |
| $\delta_{M1} = \delta_{N2}$ | Diffusion coefficient for individuals with microblades | $0.02 \text{ ky}^{-1}$. Assumed to exceed $\delta_C$ by 2 – 10 times*. |
| $r_{N2}$ | Demographic expansion rate for northerners | $43 \text{ ky}^{-1}$. Corresponds to a 4.3% annual intrinsic growth rate, slightly greater than the 3% – 4% growth rate observed for modern humans (Ammerman and Cavalli-Sforza, 2014)*. |
| $r_{M1}$ | Expansion rate for southerners with microblades | $43 \text{ ky}^{-1}$. Assumed identical to $r_{N2}$*. |
| $r_{C1}$ | Expansion rate for southerners without microblades | $28 \text{ ky}^{-1}$. Assumed to fall below $r_{M1}$*. |
| $m$ | Rate of removal from population due to malarial infections in northerners | $100 \text{ ky}^{-1}$*. The baseline value corresponds to a 1% incidence of severe malaria *per capita* when 10% of mosquitoes are infectious. |
| $b$ | Competition coefficient between the sub-populations | $5 \text{ ky}^{-1}$*. |
| $\epsilon$ | Base rate of cultural transmission between southern groups | $0.25 \text{ ky}^{-1}$. Identical to Aoki et al. (2023) and rescaled to reflect the change in units. |
| $w$ | Extent of conformity in transmission between southern populations | $0.3^{**}$. Identical to Aoki et al. (2023). |
| $L_1$ | Carrying capacity for non-microblade possessing southerners | $1^{**}$. Non-dimensionalized, so that $C_1$ represents population *density* rather than size. |
| $K_1$ | Carrying capacity for microblade possessing southerners | $1^{**}$. Assumed equal to $L_1$. |
| $K_2$ | Carrying capacity for northerners | $1^{**}$. Assumed equal to $L_1 = K_1$. |
| $c$ | Rate of cultural transmission between southern and northerners | $0.15 \text{ ky}^{-1}$*. |
| $g$ | Baseline demography rate in mosquito populations | $7000 \text{ ky}^{-1}$. Assumed from the average lifespan of an *Anopheles* mosquito (2 weeks – 2 months in cold weather), scaled by the mosquito-to-human ratio in Southeast Asia (Krajacich et al., 2020; Mehra et al., 2025) |
| $V$ | Measure of *fitness loss* for individuals without microblades at lowest temperature | $0.5^{**}$. |

(\*) denotes a parameter that requires fitting. (\*\*) denotes a dimensionless parameter.

For each simulation of System (2), the subsets of the spatiotemporal domain in which the density *M*_1_(*x, t*) is greatest are noted. Additionally, in each simulation, the time-point *t* = *T_m_* at which

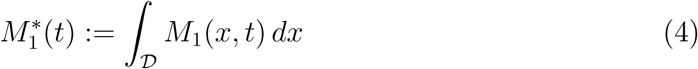

reaches a peak is recorded. In particular, *T_m_* is equivalent to the point at which the density *M*_1_ of southerners with microblades reaches its highest local maximum as a function of *t*, when integrated across the entire spatial domain. Informally, *T_m_* is referred to as the time at which microblade use by southerners is *concentrated* across the domain *D*.

For the baseline parameter values in Table 1, the subsets of *D* where *M*_1_(*x, t*) is maximal closely follow the propagating front between northerners and southerners (Figure 1). When *r_C_*_1_, the intrinsic growth rate of the microblade non-possessers, is 35% *smaller* than the corresponding rate *r_M_*_1_ = *r_N_*_2_ for microblade-possessers, the maximal value of *M*_1_ moves south linearly as *t* increases, reaching the southern end of the domain at *t ≈* 11 kya (Figure 1(a)). The front between microblade possessers and microblade non-possessers is maintained for the entirety of *T*, and similarly propagates south over time (Figure 1(b)). Each 10% change in the density *M*_1_(*x, t*) is indicated by a black contour line in Figure 1(a); lighter-colored areas in the contour plots correspond to higher densities of microblade-possessers, and darker areas correspond to lower densities. The subset of *D* where *M*_1_(*x, t*) is maximal forms a curve propagating south as *t* increases. The time at which the density *M*_1_ reaches a peak across the domain is estimated at *T_m_ ≈* 11.05 kya, and indicated with a dashed white line.

**Figure 1:**
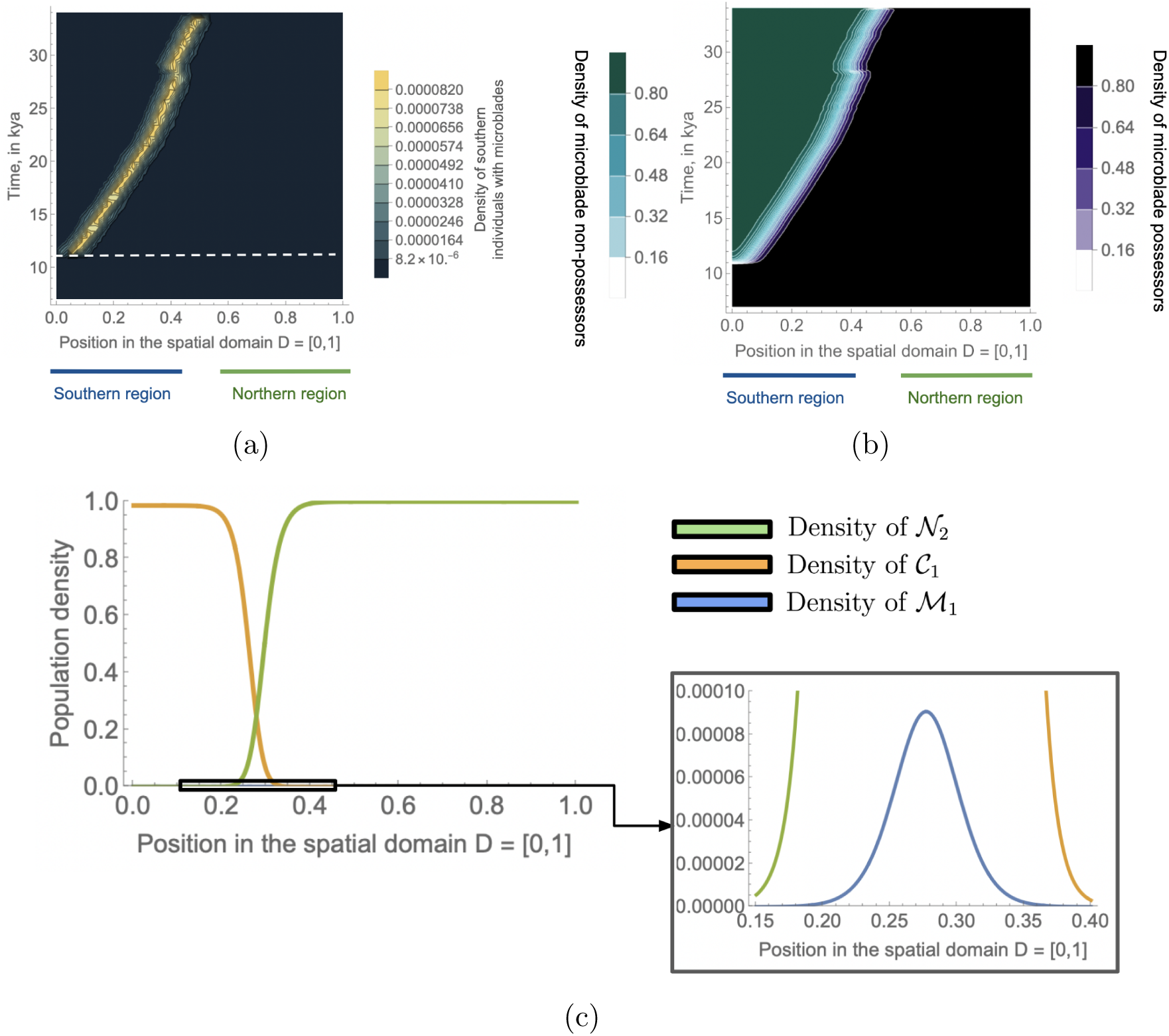
**(a)** Density *M*_1_(*x, t*) of southerners *with* microblades at each time *t ∈* [7 kya, 35 kya] and each point in *D* = [0, 1], obtained by solving system (2) numerically, when *r_C_*_1_ = 28 ky*^−^*^1^ = 0.65 *· r_M_*_1_. Values of the temperature function *f* (*x, t*) in (2) are derived from the climate data in Andersen et al. (2004), and the other parameter values are as in Table 1. Each contour line represents a 10% change in density; the brightest yellow curve enclosed by a contour represents the region of *maximal* density in the spatiotemporal domain. The southern and northern areas of the 1D spatial domain are indicated with lines underneath each panel. The dashed white line (panel A) represents the point at which *M*_1_ reaches a peak when integrated across *D*. **(b)** Total density of microblade non-possessors (*C*_1_; blue-green contours) and microblade-possessors (*M*_1_ + *N*_2_; black area with purple border) at each time-point *t*, for the same parameter values. **(c)** Density of sub-populations *C*_1_ (orange curve), *M*_1_ (blue curve), and *N*_2_ (green curve) at *t* = 20 kya. The boxed inset shows details of the region surrounding the spatial peak in *M*_1_, in the interval [0.15, 0.4] *⊂ D*.

When the diffusion coefficients *δ_C_* and *δ_M_* are increased to *δ_C_*_1_ = 0.1 ky*^−^*^1^, and *δ_M_*_1_ = *δ_N_*_2_ = 0.2 ky*^−^*^1^, the front between microblade non-possessors *C*_1_ and microblade-possessors *M*_1_ *∪ N*_2_ propagates rapidly south with time, and microblades instead concentrate at the southern border before 25 kya (Figures 2(a) and 2(c)). In contrast, when the diffusion coefficients *δ_C_*_1_ and *δ_M_*_1_ are *decreased* to *δ_C_*_1_ = 0.001 ky*^−^*^1^, and *δ_M_*_1_ = *δ_N_*_2_ = 0.002 ky*^−^*^1^, the front between *C*_1_ and *M*_1_ *∪ N*_2_ moves slowly over time (Figures 2(b) and 2(d)). In this case, the density of southerners with microblades is negligible outside of narrow regions where north-south interaction occurs, between 0.31 *≤ x ≤* 0.5.

**Figure 2:**
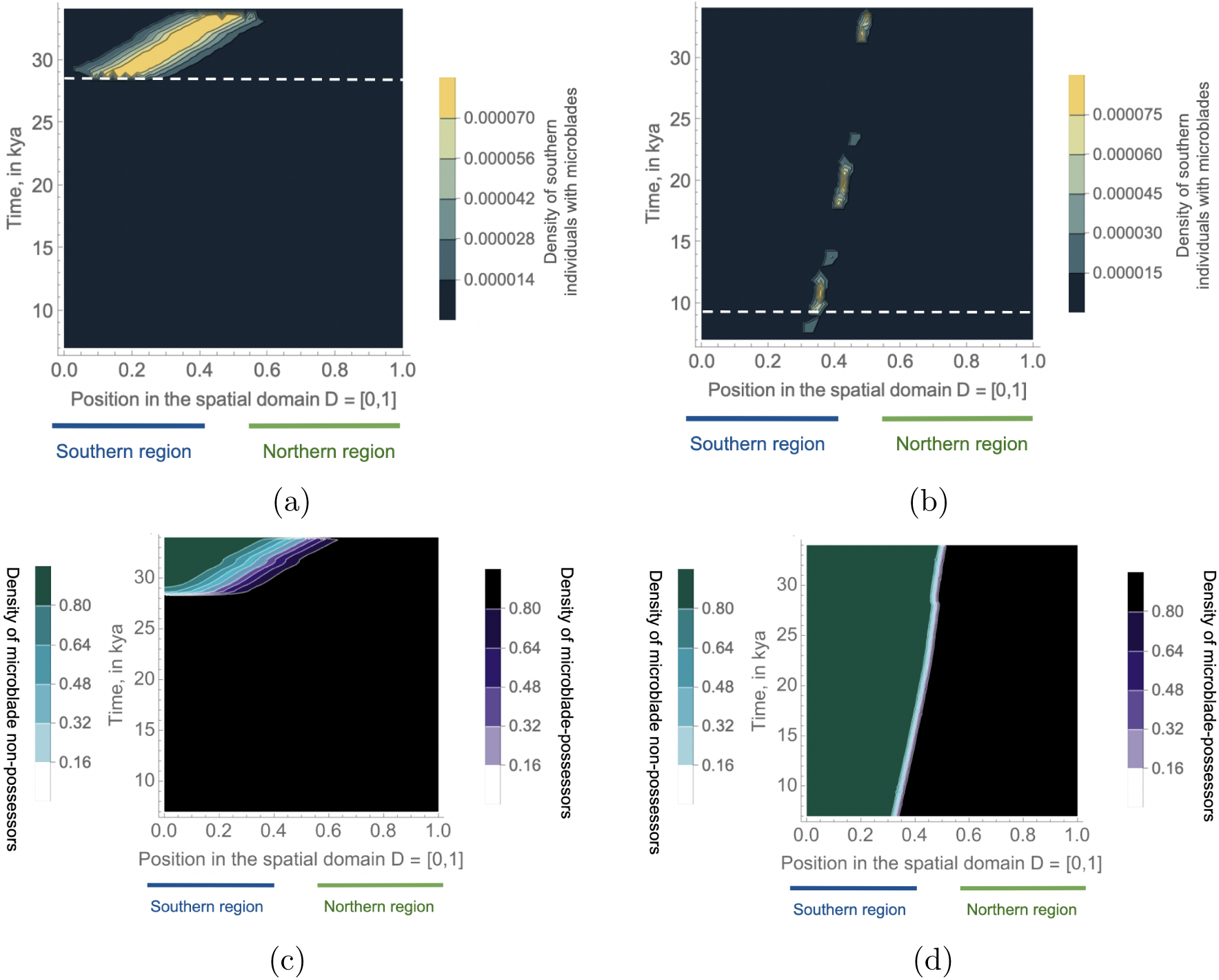
**(a)-(b)** Density *M*_1_(*x, t*) of southerners *with* microblades at each time *t ∈* [7 kya, 35 kya] and each point in *D* = [0, 1], when the microblade-assisted and microblade-free diffusion coefficients satisfy **(a)** *δ_C_*_1_ = 0.1 ky*^−^*^1^, *δ_M_*_1_ = *δ_N_*_2_ = 0.2 ky*^−^*^1^; and **(b)** *δ_C_*_1_ = 0.001 ky*^−^*^1^, *δ_M_*_1_ = *δ_N_*_2_ = 0.002 ky*^−^*^1^. Each contour line represents a 10% change in density; the bright yellow curve enclosed by a contour represents the region of *maximal* density on the spatiotemporal domain. In each panel, the dashed white line represents the point at which *M*_1_ reaches a peak when integrated across *D*. **(c)-(d)** Total density of microblade non-possessors (*C*_1_; blue-green contours) and microblade-possessors (*M*_1_ + *N*_2_; purple-black contours) at each time-point *t*, when **(c)** *δ_C_*_1_ = 0.1 ky*^−^*^1^, *δ_M_*_1_ = *δ_N_*_2_ = 0.2 ky*^−^*^1^; and **(d)** *δ_C_*_1_ = 0.001 ky*^−^*^1^, *δ_M_*_1_ = *δ_N_*_2_ = 0.002 ky*^−^*^1^. The other parameter values are as in Table 1.

The location and direction of the front vary greatly when the rate *m* of malaria transmission to northerners is varied between 30 ky*^−^*^1^ and 300 ky*^−^*^1^ (representing *per capita* incidence between 0.3% and 3%, respectively, when 10% of mosquitoes are infectious). When *m* = 30 ky*^−^*^1^, the disease burden is low enough that northerners sweep through the entire domain before 15 kya, for all values of the Lotka-Volterra competition coefficient *b >* 1.5 ky*^−^*^1^ (Figure 3(a)). As *m* increases, the location of the front at 15 kya moves *northward* through the spatial domain. When the competition coefficient *b* satisfies *b* = 2 ky*^−^*^1^, southerners reach the northern border whenever *m ≥* 110 ky*^−^*^1^. When *b* = 5 ky*^−^*^1^, significant competition prevents southerners from reaching the northern border unless *m ≥* 220 ky*^−^*^1^. The pattern of the front in Figure 1 is recovered in a diagonal region of the (*m, b*)-parameter space, in which the front propagates south over time, and lies in the interval (0, 0.5) at *t* = 15 kya (Figures 3(a) and 3(b)).

**Figure 3:**
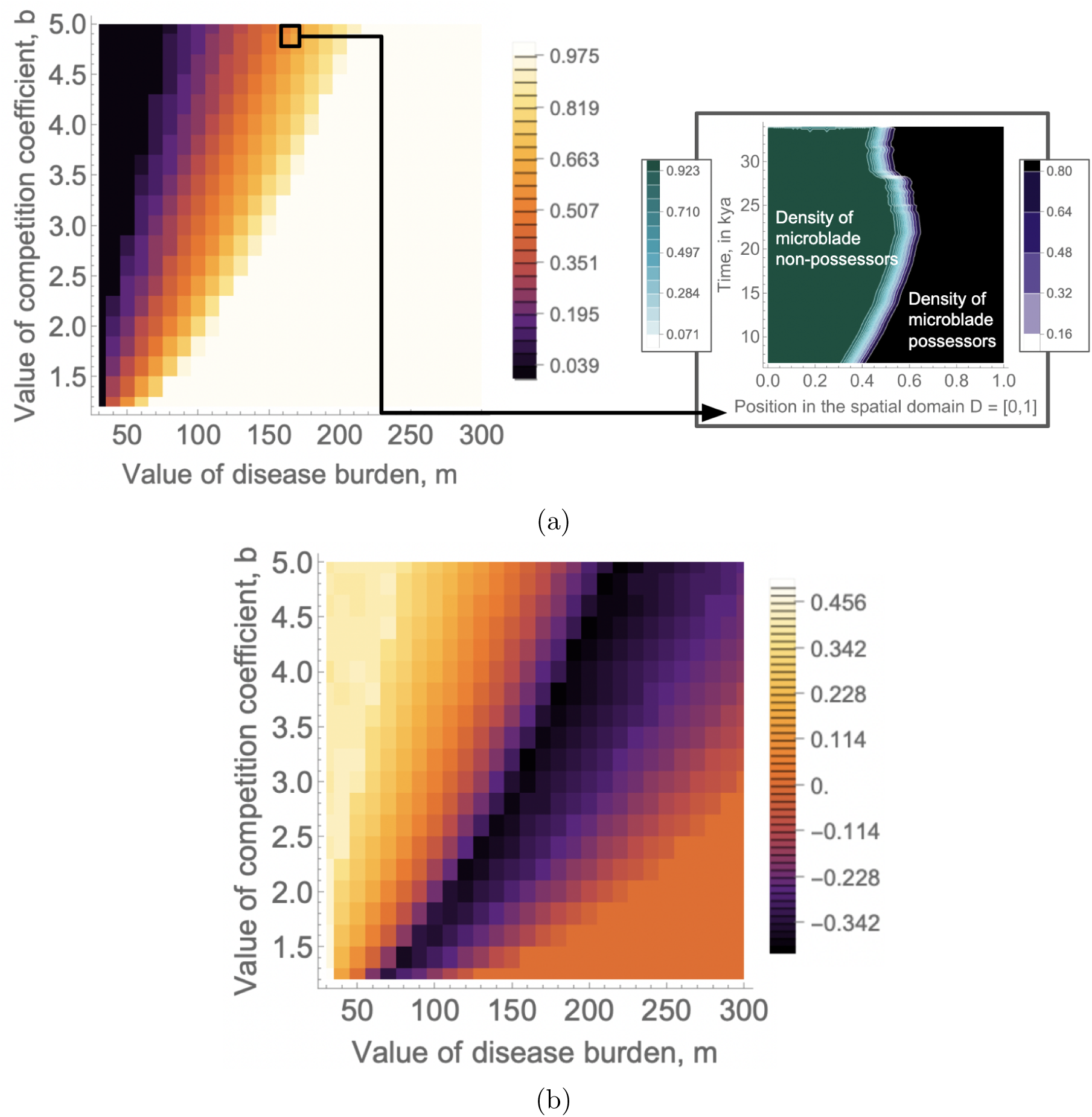
**(a)** Location of the front in *D* = [0, 1] between microblade-possessors and non-possessors, at time *t* = 15 kya, as a function of the rate *m* of severe malaria in northerners, and the Lotka-Volterra competition coefficient *b* between the three subpopulations. Darker colors indicate a more *southern* location of the front, and black indicates that microblade-possessors have reached the southern border. Similarly, lighter colors indicate a more *northern* location of the front, and white indicates that microblade non-possessors have attained the northern border. The boxed inset shows the front between microblade possessors and non-possessors for *m* = 160 ky*^−^*^1^ and *b* = 5 ky*^−^*^1^. **(b)** The *distance* that the front travels between *t*_1_ = 30 kya and *t*_2_ = 15 kya, as a function of *m* and *b*. Negative values indicate that the front has moved *north*, and positive values represent a *southward* movement of the front. The other parameter values are as in Table 1.

## 3 Existence of a stationary front between southern and northern populations before the Holocene

Throughout the Upper Paleolithic, microblades were essentially absent from archaeological findings south of the H-Q line, and the earliest recorded appearance of microblades in south China dates to 8 *−* 5 kya in the mid-Holocene (Huan et al., 2025). Moreover, population-genetic data suggests that northern and southern foragers had limited interaction up till the early Neolithic (Yang et al., 2020). These observations suggest that the front observed by Aoki et al. (2023) between northern and southerners could have remained intact before the beginning of the Holocene (11-12 kya).

Here, we estimate combinations of parameter values for which the solution of system (2) leads to a stationary front in the time *sub-interval T ^∗^* := [35 kya, 12 kya]. For greater numerical tractability, the original temperature function from Andersen et al. (2004) was first *smoothed* with a Gaussian filter (Appendix I) and separated into *N* = 100 discrete *heating* and *cooling* events. System (2) was then solved in succession for each of these discrete climate events.

In addition, the *non-spatial* component of system (2) is analyzed in order to assess when the system exhibits temporal bistability in the interval *T ^∗^* required for the existence of a stationary front. For each *i*-th climate event, modeled by a new temperature function 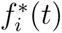 on a restricted time interval 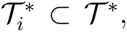 the resulting non-autonomous *_i_* ODE system takes the form

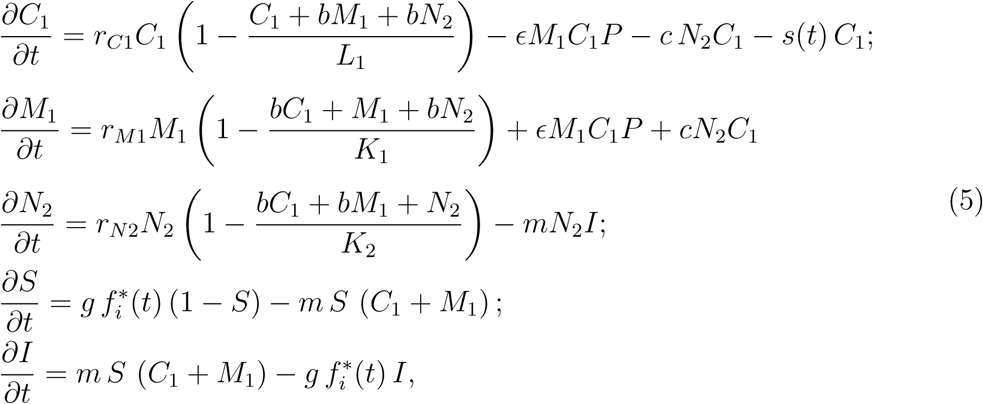

where 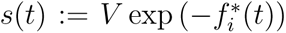 is, as before, the loss of fitness due to microblade non-possession at time *t*, and 1 *≤ i ≤* 100. System (5) is solved numerically in succession for each discrete climate event, and the resulting flows are concatenated.

For the baseline parameter values in Table 1, system (5) exhibits two bistable limiting solutions—one solution *ζ*_1_ in which *C*_1_ = 1 and *M*_1_ + *N*_2_ = 0, and another solution *ζ*_2_ in which *C*_1_ = 0 and *M*_1_+*N*_2_ = 1 (Figure 4(a)). Increasing the parameter *m* controlling the *disease burden* on northerners significantly reduces the subset of initial conditions for which the microblade-dominating solution *ζ*_2_ is stable (Figure 4(b)). On the other hand, the basins of attraction for *ζ*_1_ and *ζ*_2_ are only mildly dependent on the relative intrinsic growth rates *r_C_*_1_ and *r_M_*_1_, and on the maximum loss of fitness *V*.

**Figure 4:**
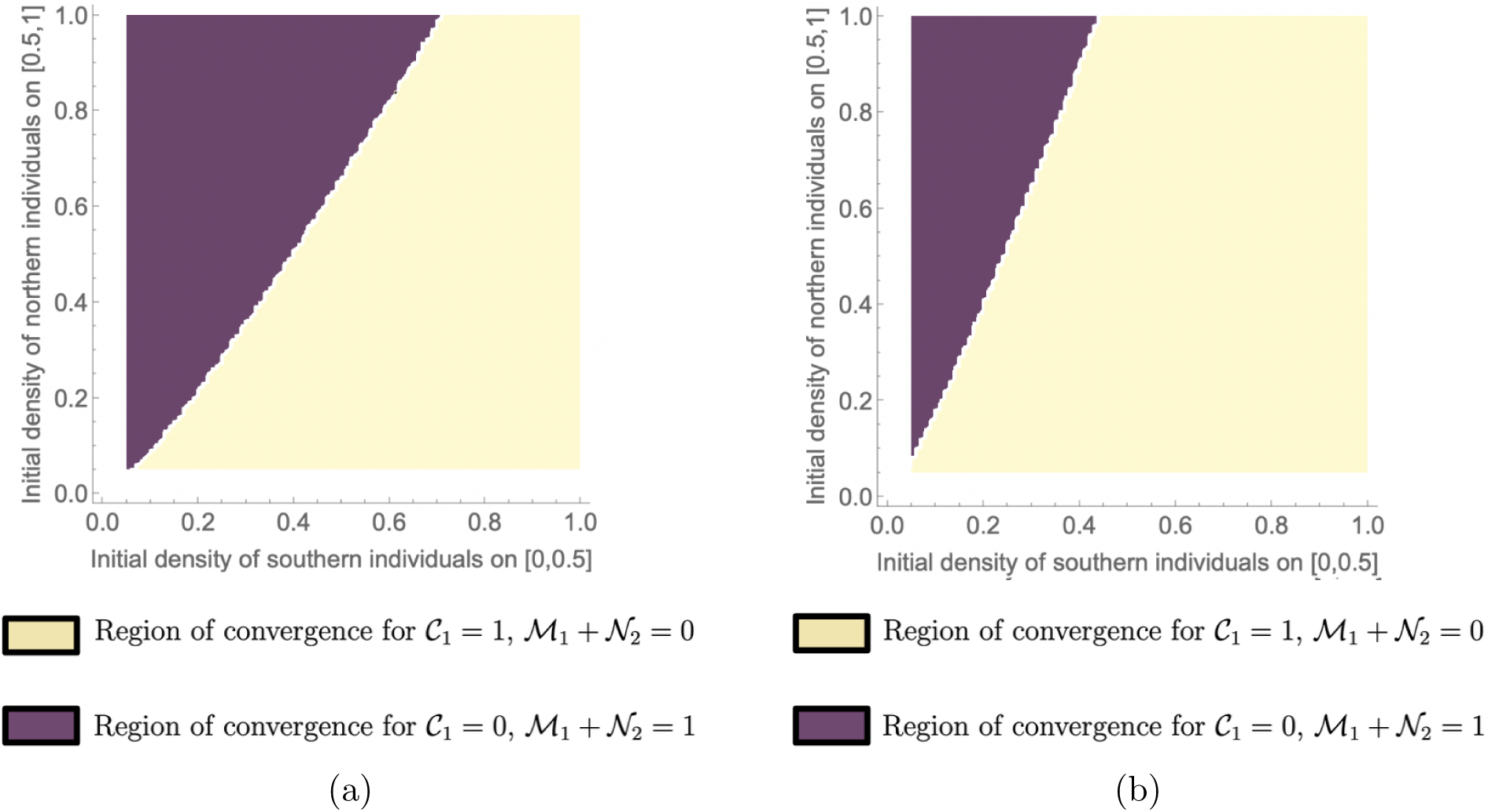
**(a)** Regions of (numerical) convergence to the two limiting solutions of system (5) for the baseline parameter values in Table 1. **(b)** Regions of convergence to the two limiting solutions when *m* = 600 ky*^−^*^1^; all other parameters are as in Table 1. In both panels, the region in which solutions converge to *N*_1_ = 1 and *M*_1_ + *N*_2_ = 0 is indicated in yellow; the region in which solutions converge to *N*_1_ = 0 and *M*_1_ + *N*_2_ = 1 is indicated in purple.

For an arbitrary temperature function 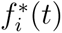 in system (5), the non-autonomous nature of the ODE system makes *analytical* conditions for bistability difficult to establish. On the other hand, when the value of the temperature function is a fixed constant, a standard local stability analysis (Appendix II) yields regions of bistability and multistability for equilibrium solutions of system (5).

In contrast to the non-spatial model, the behavior of the full spatial system in eqs. (2) over the time interval *T ^∗^* depends strongly on the values of *r_C_*_1_ and *r_M_*_1_ (Figure 5). For small values of *r_C_*_1_ and *r_M_*_1_, increases in the intrinsic growth rate of microblade-possessors relative to non-possessors results in a *southward* movement of the front (Figure 5(a)). On the other hand, for larger intrinsic growth rates such as those in Table 1, the densities of the competing sub-populations decay rapidly at the front, and high values of *r_M_*_1_ relative to *r_C_*_1_ instead result in the front propagating *north* (Figure 5(b)).

**Figure 5:**
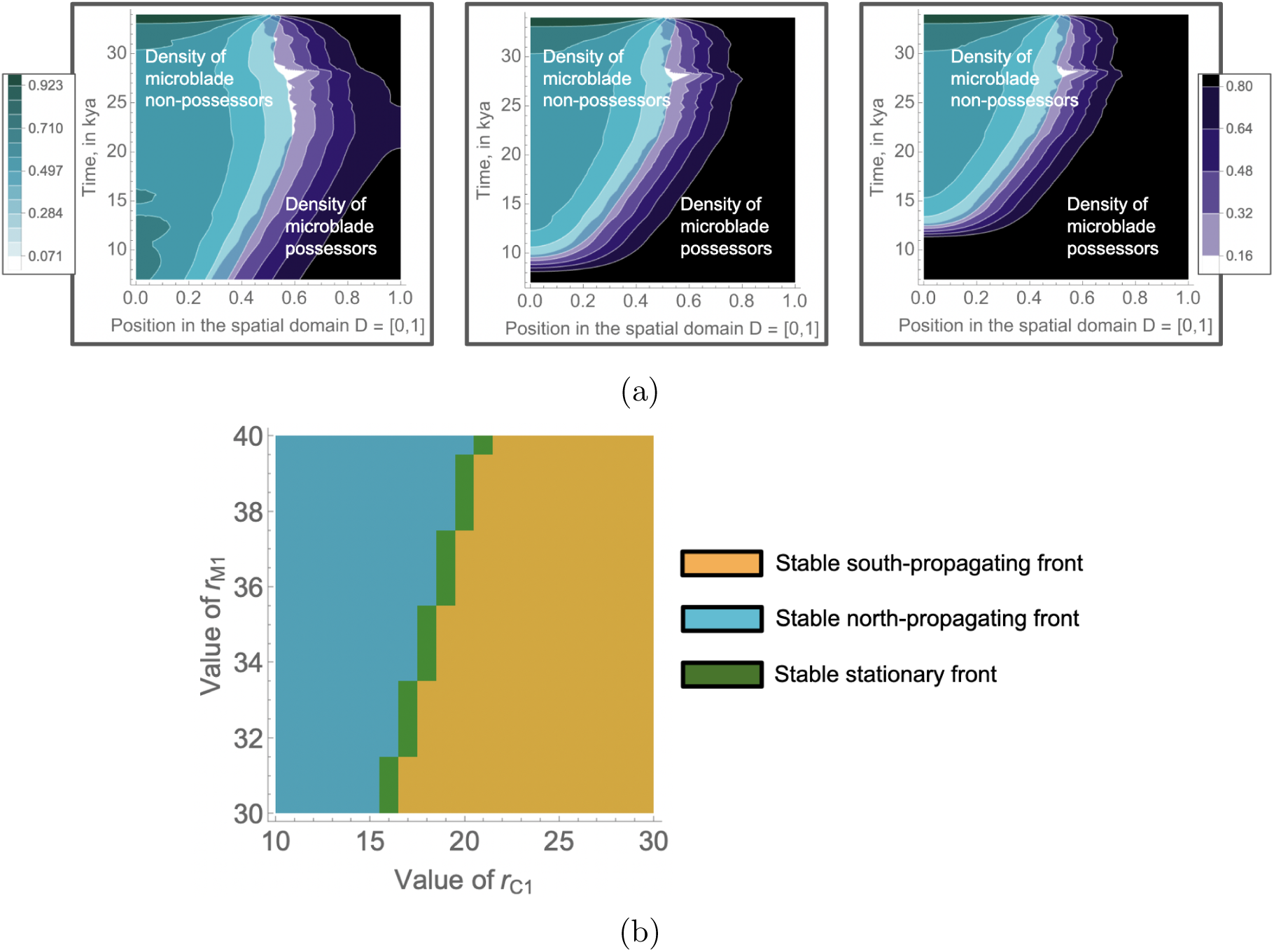
**(a)** Position of the front between microblade possessors and non-possessors, where *r_C_*_1_ = 1.1 ky*^−^*^1^ and *r_M_*_1_ = 1.1 ky*^−^*^1^ (leftmost image); *r_M_*_1_ = 2 ky*^−^*^1^ (center image); and *r_M_*_1_ = 3 ky*^−^*^1^ (rightmost image). **(b)** Stable and stationary fronts from the behavior of system (2). Values of the intrinsic growth rates are *r_C_*_1_ *∈* [10 ky*^−^*^1^, 30 ky*^−^*^1^] and *r_M_*_1_ *∈* [20 ky*^−^*^1^, 40 ky*^−^*^1^], where all other parameters are as in Table 1. Orange represents a *locally stable* south-propagating front, green a locally stable stationary front, and light blue a locally stable north-propagating front. Local stability is determined by sampling a discrete grid of initial conditions for *C*_1_*, N*_2_*, S*, and *I*; a locally stable front is assumed to be *stationary* if its location in *D* varies by less than 0.02 across the time interval *T ^∗^*.

For the parameters in Table 1, numerical simulation suggests a locally stable *north-propagating* front when *r_M_*_1_ is much larger than *r_C_*_1_, a *south-propagating* front when *r_C_*_1_ is approximately more than half of *r_M_*_1_, and a (numerically) *stationary* front when *r_C_*_1_ lies between these two regions of parameter space (Figure 5(a)). The solutions of system (2) may therefore yield a near-stationary front in *T ^∗^* even when the temperature is varied and the rate of migration is comparatively large.

## 4 Discussion

Aoki et al. (2023) proposed that the limited microblade diffusion from north to south China during the Upper Paleolithic may have resulted from the susceptibility of northerners to endemic south-based diseases, including *P. vivax* malaria or human papillomavirus. For various combinations of parameter values specifying the rates of northern contagion and population growth, their theoretical model yielded a *stationary front* between northern and southern populations, with the densities of northerners negligible on the southern side of this boundary. This could explain the archaeological finding that microblades were absent from the south on the H-Q line between the LGM and the beginning of the Holocene.

This paper investigates the disease-barrier hypothesis of Aoki et al. (2023) for microblade diffusion in the context of the sharp *climate changes* characterizing the Upper Paleolithic and early Holocene. It is assumed in particular that one of the pathogens driving excess disease burden for northerners below the Qinling–Huaihe (H-Q) Line was *climate-dependent*, and in particular vector-borne (such as *P. vivax*, transmitted by *Anopheles* mosquitoes (Gething et al., 2011)). We then model the effect of distinct climate events on the front between the northern and southern populations.

Our extended model in Section 2 simulates how the *climate history* of the Upper Paleolithic could have influenced the time at which southerners converted to microblade use. This extended model incorporates two directions for cultural transmission of microblade use—*external*, from northerners to southerners, at the front where they interact; and *internal*, from southerners who possess microblades to southerners who do not. Projections of temperature between 35 and 7 kya, obtained by the North Greenland Ice Core Project (NGRIP; Andersen et al. (2004)), are then used to simulate the times at which microblade use *peaks* among southern populations.

The times at which microblade use is concentrated across the southern population is sensitive to the migration rates *δ_M_*_1_ and *δ_C_*_1_ of microblade possessing and non-possessing individuals, respectively, and to the extent of disease burden *m* on the northern population. When *δ_M_*_1_ = 0.02 ky*^−^*^1^ and *δ_C_*_1_ = 0.01 ky*^−^*^1^ (equivalent to a mean annual displacement of 0.47 km and 0.33 km, respectively) and *m* is set to 100 ky*^−^*^1^ (equivalent to an 1% annual incidence of exit-causing malaria when 10% of mosquitoes are infectious), microblade use reaches the southern edge of the domain between 20 and 10 kya (Figure 1), after temperatures have begun to rise following the LGM.

However, when the extent of disease burden *m* is sufficiently high (e.g., *m* = 160 ky*^−^*^1^, equivalent to an 1.6% annual incidence of severe malaria when 10% of mosquitoes are infectious), southerners initially *expand* into northern territory before eventually withdrawing from the north after 21 kya (Figure 3, boxed inset). This withdrawal may have resulted from a decrease in the estimated temperature between 23 and 20 kya (Figure 6, blue curves), which could have temporarily *lowered* the high simulated disease burden on northerners. As a result, microblades use moves into southern territory, but never reaches its southern border.

When the migration rate is especially low (corresponding to low values of the diffusion coefficients *δ_C_*_1_*, δ_M_*_1_*, δ_N_*_2_), the propagation of the front becomes particularly slow, approaching the stationary front in Aoki et al. (2023). In this case, southern microblade use remains limited to a narrow *interaction* region at the location of the front (Figures 2(b) and 2(d)). Moreover, at moderate values of the diffusion coefficients *δ_C_*_1_*, δ_M_*_1_*, δ_N_*_2_, a two-dimensional set of intrinsic growth rates (*r_C_*_1_*, r_M_*_1_) results in the formation of a locally stable stationary front (Figure 5(b)).

The dependence of northern migration on temperature change could suggest a possible hypothesis for why microblades reach the south only several millennia following the Last Glacial Maximum, in the early-to-mid Holocene. In the cold climate of the LGM, decreases in the transmission of vector-borne disease could have allowed northerners to encroach into southern territory, resulting in an initial population of southerners who possessed microblades. Then, if the rise of temperatures following the LGM caused the northerners to *withdraw* from southern territory, competition for resources could have occurred primarily between southerners *with* microblades and those without them— potentially resulting in cultural conversion to microblade use.

We briefly discuss some limitations of these model extensions below. First, we do not consider the hypnozoite-induced *relapses* that distinguish *P. vivax* malaria, and could affect northerner disease burden even after departure from southern territory. Additionally, our models do not incorporate the potentially complex effects of climate on migration rates in the Upper Paleolithic. In particular, the onset of the LGM increased emphasis on hunting, which likely spurred additional migration (Spikins, 1997), but is simultaneously associated with geographical barriers such as ice sheets that could hinder travel (Liu et al., 2020).

Although some of the model parameters we use are sourced from existing literature (i.e., the diffusion rate of microblade non-possessers), the majority of the parameter values used here have limited empirical evidence. Future analyses of the time at which microblades first reached the south should include a detailed sensitivity analysis, to understand the spatial dynamics of microblade transmission across the full range of parameter space.

Furthermore, the two separate directions of cultural transmission in system (2)— between northern and southerners, and within the southern population—motivate the problem of estimating the *relative contributions* of these directions to the eventual density of microblades in the south. Our numerical simulations (Figures 1–3) suggest that the model represents *demic* diffusion of microblade use, in which the pattern of microblade use follows the spread of northerners into southern territory, and most transmission occurs at the front between northerners and southerners (Cavalli-Sforza et al., 1993). Extensions of this study could study the conditions under which *internal* cultural transmission contributes non-negligibly to microblade uptake in the south.

The role of climate in directing migration and human behavior throughout Upper Paleolithic Asia has been subject to substantial study ((Soares et al., 2008; Dong et al., 2020)), but to our knowledge has not been applied to the diffusion of microblades across the Qinling-Huaihe line. Our analysis suggests that analysis of the role of climate on cultural transmission could have broader applications in studies of cultural evolution.

## Acknowledgments

This research was supported in part by the Stanford Center for Computational, Evolutionary, and Human Genomics, and by the Morrison Institute for Population and Resource Studies at Stanford University.

## Appendix I: The non-spatial temperature function *f*^∗^(*t*)

**Figure 6:**
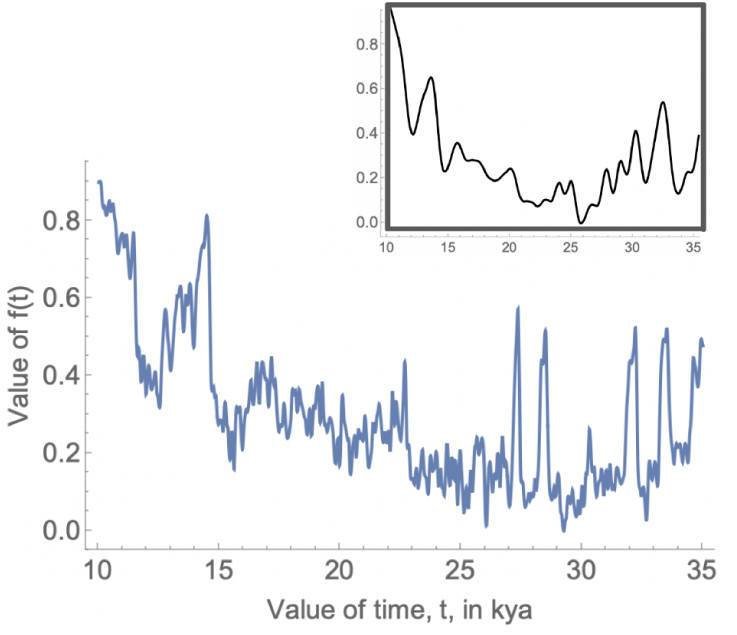
Smoothing of the non-spatial temperature function *f ^∗^*(*t*) obtained from NGRIP data in Andersen et al. (2004). The original normalized temperature function is plotted in blue; the smoothed version, obtained by convolving *f ^∗^*(*t*) with a Gaussian kernel with radius *R* = 5 years, is plotted above in black.

## Appendix II: Multistability analysis of the extended system without temperature variation

In this section, we analyze the stability of equilibria for the *non-spatial* model of microblade transmission (eqs. (5)), which in turn represents a necessary condition for the formation of a stationary front between microblade-possessors and non-possessors. The analysis here is limited to the simple case in which the non-spatial temperature function *f^∗^*(*t*) is *constant* for all *t*.

In particular, fixing *f^∗^*(*t*) at a constant *F ∈* R^+^, the non-spatial system associated with system (2) becomes

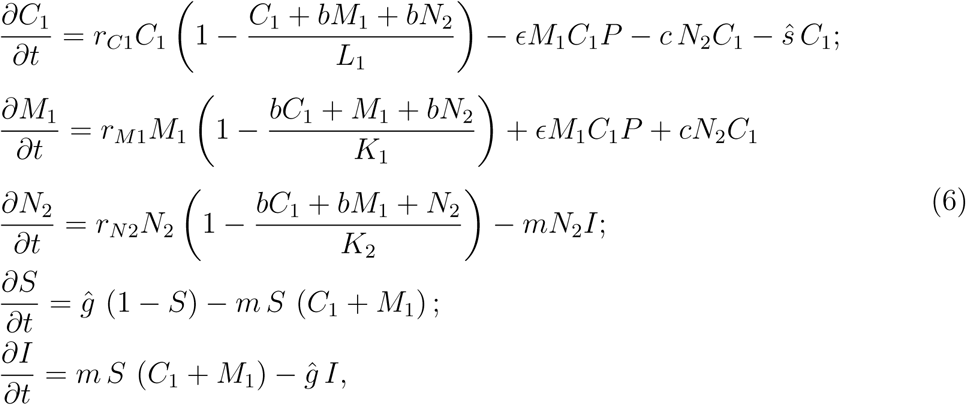

where *ĝ* := *ĝ · F* and *ŝ* := *V e^−F^*. We begin by considering all equilibrium solutions for system (6) in which exactly two of the three human populations have zero density. This yields three non-extinction equilibria of the form

- The *northerners-only* equilibrium (*ζ*_1_):

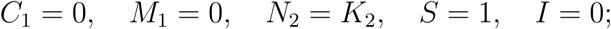
- The southern *microblades-only* equilibrium (*ζ*_2_):

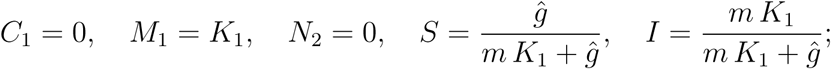
- The *no-microblades* equilibrium (*ζ*_3_):

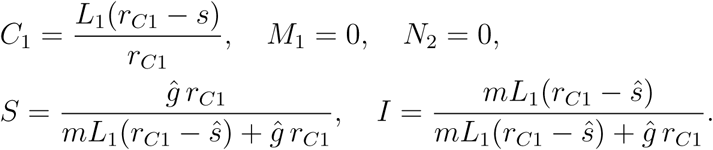

Note in particular that the no-microblades equilibrium *ζ*_3_ is biologically realistic only when the intrinsic growth rate of the microblade non-possessing southerners, *r_C_*_1_, exceeds the (constant) loss of fitness *ŝ* due to temperature. The cultural transmission term *ɛM*_1_*C*_1_*P* = *ɛM*_1_*C*_1_(*M*_1_*/*(*M*_1_+*C*_1_)*−w*) in eqs. (6) is set to 0 at the northerners-only equilibrium *ζ*_1_.

The Jacobian matrix associated with system (6) takes the form

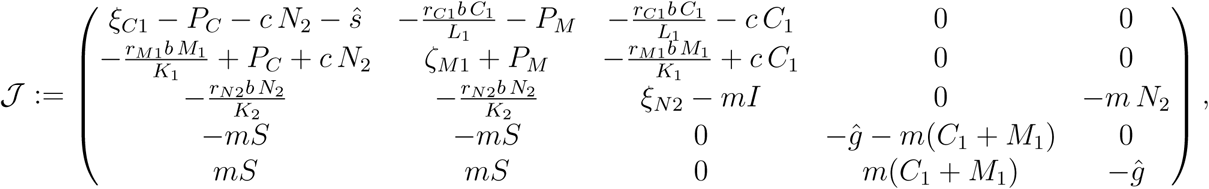

where

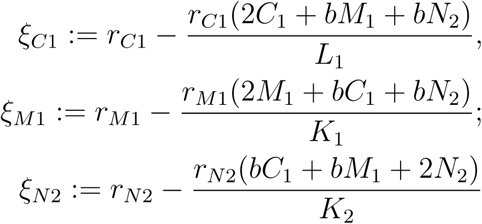

and

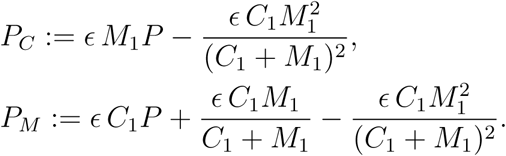

To simplify the computation below, we assume as in Aoki et al. (2023) that the three carrying capacities *L*_1_*, K*_1_*, K*_2_ are all *equal* to a constant *Q ∈* R^+^. Evaluating the Jacobian *J* at the three equilibria {*ζ*_1_*, ζ*_2_*, ζ*_3_} then yields that

i. The northerners-only equilibrium *ζ*_1_ is locally asymptotically stable (LAS) when *b >* 1;
ii. The southern microblades-only equilibrium *ζ*_2_ is LAS when *b >* 1;
iii. The no-microblades equilibrium *ζ*_3_ is LAS when *r_C_*_1_ *> s* and *b*(*r_C_*_1_ *− s*) *> r_C_*_1_.

Combining these results yields the following set of conditions for the *multistability* of equilibria in the non-spatial system (6):

### Proposition 2.1.

*When b >* 1*, the equilibria ζ*_1_ *(where N*_2_ *is the only non-empty population) and ζ*_2_ *(where M*_1_ *is the only non-empty population) are both LAS. Moreover, when r_C_*_1_ *> s and the stronger condition*

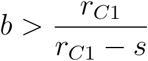

*is met, all three equilibria ζ*_1_*, ζ*_2_*, and ζ*_3_ *(where C*_1_ *is the only non-empty population) are simultaneously LAS*.

Note that the stability of the equilibria *ζ*_1_*, ζ*_2_*, ζ*_3_ does not require restrictions on the rates of cultural transmission *c, ɛ*, nor on the cultural conformity parameter *w*.

